# Survival of tardigrades (*Hypsibius exemplaris*) to subzero temperatures depends on exposure intensity, duration, and ice-nucleation—as shown by large-scale mortality dye-based assays

**DOI:** 10.1101/2024.02.28.582259

**Authors:** Ana M Lyons, Kevin T Roberts, Caroline M Williams

## Abstract

Tardigrades are an emerging model system for understanding a diversity of environmental stress responses, yet few studies describe the physiology of cold tolerance in hydrated, active tardigrades. Here, we develop methods to screen tardigrades for survival in a high-throughput manner, to investigate the impacts of several key environmental conditions on survival. The visualization of dye uptake (SYTOX Green) in hydrated, cold-exposed *Hypsibius exemplaris* allows us to quickly and accurately quantify the survival of thousands of animals, under a range of ecologically-relevant low temperatures, exposure times, conditions, and thermal acclimations. As a proof-of-concept, we show that SYTOX Green uptake more accurately predicts 2-week survival outcomes of tardigrades post-cold exposure, compared to previous methods of scoring survival (locomotion). We show that hydrated, active tardigrades survive mild cold exposures of - 10°C at high rates of ∼98%. Survival of tardigrades to exposures of -15°C depends on environmental freezing in pure mineral water, and survival decreased exponentially with exposure time at -20°C (to 45% after 24 hours; with freezing occurring at nearly all -20°C timepoints). To investigate the role of environmental ice-formation on tardigrade survival vs. temperature, we incubated unacclimated tardigrades with ice-nucleating bacteria—which initiate environmental freezing at higher temperatures (-1.8 to 3.8°C). Surprisingly, we found a significant increase in survival of tardigrades frozen at -20°C (p-value = 0.0152) with the addition of *Pseudomonas syringae* compared to non-inoculated controls, as well as observing high-survival of tardigrades in ice-nucleated samples exposed to -10°C and -15°C. This indicates the species’ tolerance to environmental ice formation and exposure to our lowest temperature (-20°C), under certain conditions of controlled environmental ice formation. A 3-week acclimation of tardigrades to mild cold (1°C and 4°C) in constant darkness did not significantly improve survival after acute exposure to low temperature, but acclimating animals to 15°C did. Overall, we find that *H. exemplaris*—an emerging tardigrade model species—has a range of cold tolerance capabilities, dependent on time, temperature, environmental ice-formation, and culturing conditions. This work offers a framework with new tools for performing large-scale physiological assays in numerous species, establishing tardigrades as a tractable and uniquely informative model system in comparative physiology and the study of environmental stress.

**Key Findings:**

- Active, hydrated tardigrades—with no prior acclimation to cold—have high survival of temperatures above -15°C, even in response to prolonged exposures.
- Below -15°C, tardigrade survival declines exponentially with increasing exposure time.
- Incubating tardigrades with ice-nucleating bacteria significantly improves survival after cold exposure, illustrating the importance of ice-formation dynamics and environmental microbes.
- A 3-week acclimation of tardigrades to mild cold (1°C & 4°C) does not significantly improve survival to low temperature, while acclimation to 15°C (vs the standard culture condition of 20°C) does.
- Uptake of the dye SYTOX Green is a more accurate metric of tardigrade mortality in response to cold exposure, compared to the traditional method of scoring lack of locomotion during recovery.

## Introduction

Tardigrades are a group of microscopic animals that have captured the curiosity of scientists, both for their abilities to survive environmental extremes, and for their high potential to be developed as a model system for a wide range of biological questions (Goldstein 2018) and translational applications (Giovannini et al. 2022; Piszkiewicz et al. 2019; Kirke et al. 2020; Hashimoto et al. 2016; Schill et al. 2009). Under certain environmental stresses, tardigrades evade harm by entering various forms of protective dormancy known as cryptobiosis or encystment, as seen in desiccation, osmotic stress, or radiation (Møbjerg and Neves 2021). As one especially striking example of survival, researchers found that a species of Antarctic tardigrades *Acutuncus antarcticus* could be successfully revived from storage in the tun state (desiccated) at -30°C for over 30 years, with no addition of cryoprotectants or special storage protocols. These forms of protective dormancy involving tun or cyst formation, however, are not often seen in cases of cold-exposed tardigrades originating from less-extreme habitats (Guidetti and Møbjerg 2018; Guidetti et al. 2011).

Instead, many species of cold-exposed tardigrades are believed to go into a physiological state known as cryobiosis, where active animals can survive low temperatures and their overall morphology remains largely unchanged, compared to tun or cyst formation (Hengherr and Schill 2018). Supporting this claim, a recent ecological survey of microfauna in snow algae blooms in northern Japan indicated that physiologically active tardigrades were among the most abundant organisms, thus suggesting that tardigrades do not rely on deep dormancy in snowy environments (Ono et al. 2021). Likewise, a microbiome diversity survey conducted in British Columbia detected a high abundance of snow algae genetic material in tardigrade gut contents, indicating snow algae consumption by tardigrades during winter months (Yakimovich et al. 2020). Despite these observations indicating tardigrades’ abilities to tolerate winter conditions, few studies have rigorously characterized the impact of cold on hydrated, active tardigrades, especially under ecologically relevant conditions where exposure time, temperature, and environmental ice-inoculators may vary (Schill 2018). Methodological challenges in handling and scoring these unique microscopic animals—especially with the high accuracy and throughput needed to interrogate interactions between these crucial factors—have historically contributed to this gap in knowledge.

Low temperatures and freezing are undoubtedly a major selective pressure that tardigrades experience, given their global distribution in cold climates with seasonal or frequent subzero temperatures (McInnes and Pugh 2018). All tardigrade species require a film of liquid water in order to locomote, feed, and reproduce, and ice-formation threatens this requirement, presumably impacting essential elements of tardigrades’ metabolic function, cellular integrity, and other temperature-dependent physiological processes (Bertolani et al. 2004). As microscopic water-dwelling organisms, tardigrades are especially vulnerable to rapid shifts in ice-formation, yet environmental factors impacting their survival to subzero temperatures and ice-formation are not well understood. In order to survive subzero temperatures while hydrated, tardigrades are believed to deploy one of two major cold tolerance strategies: freeze avoidance or freeze tolerance (Hengherr and Schill 2018). Organisms using freeze-avoidant strategies cannot survive internal ice formation, and use colligative solutes such as glycerol or trehalose to suppress the freezing point of their body fluids or employ cryoprotective dehydration to avoid ice formation (Denlinger and Lee 2010). Freeze tolerant organisms can survive extracellular or even intracellular ice formation, and often do so by inoculating ice formation at relatively warm subzero temperatures using ice-nucleating proteins, resulting in ice formation occurring more slowly (Denlinger and Lee 2010). It remains unknown if tardigrades produce their own antifreeze or ice-nucleating proteins, or if they are influenced by the ice-nucleating proteins or agents from any neighboring microorganisms, such as bacteria or fungi that share their winter-time habitat.

Although several studies have attempted to describe the cold tolerance strategies of tardigrades using calorimetry, with the goal to detect liquid to ice phase transitions in tardigrade samples, conclusions of the exact mode, plasticity, and robustness of cold tolerance remain ambiguous (Westh et al. 1991; Westh and Kristensen 1992). Early calorimetric studies conducted on limno-terrestrial species *Richtersius* (*Adorybiotus*) *coronifer* and *Amphibolus nebulosus* detected relatively high ice-crystallization temperatures (-6 to -7°C) for tardigrades, suggesting freeze tolerance as the primary strategy to survive cold (Westh and Kristensen 1992). Likewise, these authors predicted that ice nucleators likely played an important role in modulating the higher-than-expected freezing temperatures for these tardigrade species (Westh et al. 1991). More recent studies on nine tardigrade species indicated that phase transitions occurred at much lower temperatures, with tardigrades experiencing supercooling points ranging from roughly - 10°C to -24°C, although most species survived ice formation well (Hengherr et al. 2009). However, most prior laboratory conditions lack ecological relevance, being carried out in minimal water volumes, on starved animals, and without the presence of any environmental microbes. We thus aimed to assess the impact of more environmentally relevant conditions on the cold tolerance strategy.

Here, we develop a series of phenotypic assays in the cosmopolitan eutardigrade species *Hypsibius exemplaris*, to explore foundational questions in tardigrade cold tolerance. *H. exemplaris* is an emerging model system due to its ease of propagation in laboratory settings, genomic resources, and the growing collection of knowledge surrounding the species’ development, physiology, and stress tolerance (Goldstein 2018). Although several recent studies have focused on the desiccation tolerance of *H. exemplaris* and the evolution of its genome (Yoshida et al. 2017), documentation is absent on this species’ cold tolerance strategy, as well as how cold stress impacts the physiology and viability of this experimentally important species. The type specimen of *H. exemplaris* was collected from a benthic sample from a pond in Darcy Lever, Bolton, Lancashire, England (latitude 53.566166, longitude -2.3925272; British National Grid Ref. SD741078) (McNuff 2018) where animals likely experience fluctuating subzero temperatures throughout winter months, although members of this genus are believed to be globally widespread across temperate climates (GĄsiorek et al. 2018). A closely related species, *Hypsibius dujardini*, was found to have only “moderate” tolerance to cold compared to a set of six other tardigrade species studied (Guidetti et al. 2011), but whether *H. exemplaris* is more robustly cold-tolerant remains unknown. It is unclear whether *Hypsibius* genus tardigrades (and other tardigrade species) have a fixed level of cold tolerance, or if their sensitivity to subzero temperatures is dependent on fluctuations in environmental conditions such as the intensity and duration of low temperature, or the presence of external ice-nucleators. Filling this gap in knowledge would not only propel our understanding of a key aspect of this *H. exemplaris’* ecophysiology, but it would provide a more holistic context for understanding other forms of stress tolerance than this emerging model. More so, methodology developed in the more tractable *H. exemplaris* species could serve as a future framework to characterize cold tolerance phenotypes in a wide range of tardigrade species.

One limitation of previous studies assessing tardigrade cold tolerance (or any physiological assay that requires scoring tardigrade mortality) is the difficulty in determining whether animals are dead or alive, based on locomotion alone. For example, the largest cold tolerance study of tardigrades to date indicated that fully hydrated *Ramazzottius varieornatus* exhibit high levels of cold tolerance after exposures to -20°C, -80°C and -196 °C, but the scoring of animals relied on manual visualization of locomotion alone, which is a laborious and time-sensitive process, limiting throughput (Møbjerg et al. 2022). Additionally, lack of locomotor activity can indicate mortality but can also indicate dormancy or temporary lack of motivation or ability to move (e.g. due to molting cycles), introducing considerable noise into survival measurements. Furthermore, tardigrades’ microscopic size and transparent color can make such behavioral observations difficult and irreproducible. Thus, we optimized protocols to stain tardigrades with the live/dead cell marker dye SYTOX Green. SYTOX Green is a commonly used dye in flow cytometry to indicate mortality in high-throughput cellular screens, and it has been repeatedly demonstrated to bind to DNA only in nuclei that are highly permeabilized due to cell death (Thakur et al. 2015; Lebaron et al. 1998; Roth et al. 1997). This molecular dye was first demonstrated for use in tardigrades with a series of proof-of-concept experiments using sodium azide to induce mortality (Richaud and Galas 2018). Using this new methodological tool, we utilize tardigrades’ microscopic size and transparent color to ultimately score mortality with greater accuracy and efficiency.

Here, we use *H. exemplaris* as a model system to experimentally assess the plasticity of tardigrade cold tolerance in response to different ecologically relevant temperatures, exposure times, and key environmental conditions (the presence of external ice-inoculators and the process of thermal acclimation). In our first experiment, we validate our new method to phenotypically screen the mortality of tardigrades after cold exposure in a more accurate and high-throughput manner, using SYTOX Green dye vs. locomotion as metrics of mortality. We then dissect the combined impacts of temperature and duration of cold exposure on survival, in thousands of tardigrades, using SYTOX Green to indicate mortality. Third, we explored the impact of natural (bacterial) ice-inoculators and the role of ice-formation on tardigrade survival. Lastly, we investigated the role of thermal acclimation prior to cold exposure on survival. Overall, this work presents the first experimental evidence that tardigrade survival is significantly impacted by both the degree of low temperature and duration of cold exposure, as well as other environmental factors such as the presence of ice-inoculating bacteria and the process of acute thermal acclimation. Furthermore, methodological advancements of using SYTOX Green to score the survival of thousands of tardigrades allowed us to determine a range of scenarios where tardigrade cold tolerance varied between high (surviving freezing conditions) and low survival.

## Methods

### Tardigrade culture

Experiments were conducted on the eutardigrade species *Hypsibius exemplaris* Z151 (reclassified from *Hypsibius dujardini* in 2017), purchased from Sciento (Manchester, United Kingdom). Tardigrades were cultured in 100 x 25 mm deep-well petri dishes (USA Scientific, 8609-0625) in constant lowlight at 20°C, and animals were suspended in roughly 200mL of commercially sourced mineral water (Crystal Geyser). Tardigrades were fed with 15mL of dense *Chlorococcum sp*. (cultured from Sciento, Manchester, United Kingdom), and added bi-monthly directly to deep well petri dishes. The top 100mL of water and culture debris was skimmed from each petri dish bimonthly, via a 50mL serological pipette and pipette controller, and 100mL of fresh mineral water was added along with algae food source. Dishes were manually inspected for fungal infection (20X magnification) after each feeding, and each tardigrade culture plate was split into 2-5 new culture plates every 4 months.

To generate the tardigrade food source, *Chlorococcum sp*. algae starter cultures were added to 800mL of commercially sourced mineral water and Bold Modified Basal Freshwater Nutrient Solution (Sigma Aldrich B5282-500ML). These cultures were grown to high density (typically ∼2-3 weeks at 20°C) in parafilm-sealed, autoclaved 1000mL Corning Pyrex Erlenmeyer Flasks, under white LED light and constant aeration. Algae aeration was achieved by inserting a sterile serological pipette into the flask, which was connected to an aquarium pump (Tetra Whisper, 20-40 gallon) with plastic tubing and a 0.2μm pore syringe filter.

### Validation of SYTOX Green uptake as an improved survival assay

To determine the most accurate survival scoring metric (lack of locomotion or SYTOX Green uptake) to describe mortality in tardigrades after cold exposure, groups of tardigrades were exposed to low temperatures, initially scored with both metrics, and tracked for activity over a 14-day period (as a proxy for long-term survival outcomes). To achieve this experimental goal, groups of mixed staged adult *Hypsibius exemplaris* (n**≈**20 animals) were collected from culture dishes (at 20°C) via manual microaspiration (a Sigma-Aldrich A517 aspirator tube assembly with a hand-pulled 150mm Glass Pasteur Pipette tip, connected with ∼3cm of 1/4-Inch PVC tubing), washed in mineral water thrice to remove algae sediments in secondary petri dishes, and collected into a 1.5mL Eppendorf tubes with 100μl of mineral water (Crystal Geyser) to constitute one experimental sample. Samples (N = 10 in total, N = 2 per cold exposure time) were then exposed to cold treatments of -15°C for up to 48 hours (in increments of 4, 8, 12, or 48 hours) in a cooling circulator (VWR Scientific 1167P Heating/Cooling Recirculating Water Bath, filled with 1 part Ethylene Glycol: 1 part Water, via placement into a secondary metal container submerged into glycerol antifreeze). After the desired length of cold exposure time, cooled tardigrade samples were then removed from the cooling circulator onto the benchtop at 20°C and allowed to warm for 20 minutes. A parallel set of tardigrade samples (N = 4, with N = 1 exposed to 4, 8, 12 or 48 hours of -15°C) where not cooled and acted as a room temperature control.

Tardigrade samples were then stained with an optimized concentration (1μM) of SYTOX Green for 1 hour (according to Richaud and Galas 2018), spun at low speed on a desktop centrifuge, washed with distilled water, placed onto wet mounts without coverslips, and immediately viewed under a fluorescent dissecting microscope at 120X magnification, using a Leica DMRXA2 upright fluorescent microscope equipped with a Leica I3 long-pass GFP filter and a Leica DC500 camera (Leica Microsystems, Wetzlar, Germany). Each image was captured by using identical settings. All animals were manually scored as 1) having the presence or absence of SYTOX Green uptake and 2) having the presence or absence of locomotor activity, by visual inspection. Select animals were photographed, to illustrate representative staining patterns of SYTOX Green uptake throughout the tardigrades’ bodies. Individual animals were then transferred into wells of 48-well plates, containing 300μl of distilled water and 100μl of *Chlorococcum sp* food source. Locomotion was assessed daily for 14 days, via manual observation under 30X on a dissecting microscope, in order to determine long-term survival. Animals were scored as being “true positives” for mortality if they were observed to have coordinated locomotion for 1 day or less, during the 14-day observation period. Animals were scored as being “true negatives” for mortality if they were observed to have coordinated locomotion for at least 2 days during the 14-day observation period.

To determine which method of mortality scoring was more accurate for downstream use, we analyzed data using an Exact McNemar test (with central confidence intervals) using the exact2x2 package in R (Version 4.1.3). We conducted this analysis for all cold-exposed tardigrades in aggregate (Figure 1G), as well as when partitioned according to water-state (Supplementary Figure 1A & 1B, as well as exposure time Supplementary Figure 2). For each mortality scoring metric, specificity was defined as TN / (TN+FP), sensitivity was defined as TP / (TP+FN), and accuracy was defined as (TP + TN) / (TP + TN + FP + FN). TP = # of tardigrades that were predicted dead and were confirmed dead (among the 14 day observation period). TN = # of tardigrades that were predicted alive and were confirmed alive. FP = # of tardigrades that were predicted dead and were confirmed alive. FN = # of tardigrades that were predicted alive and were confirmed dead.

**Figure 1.**
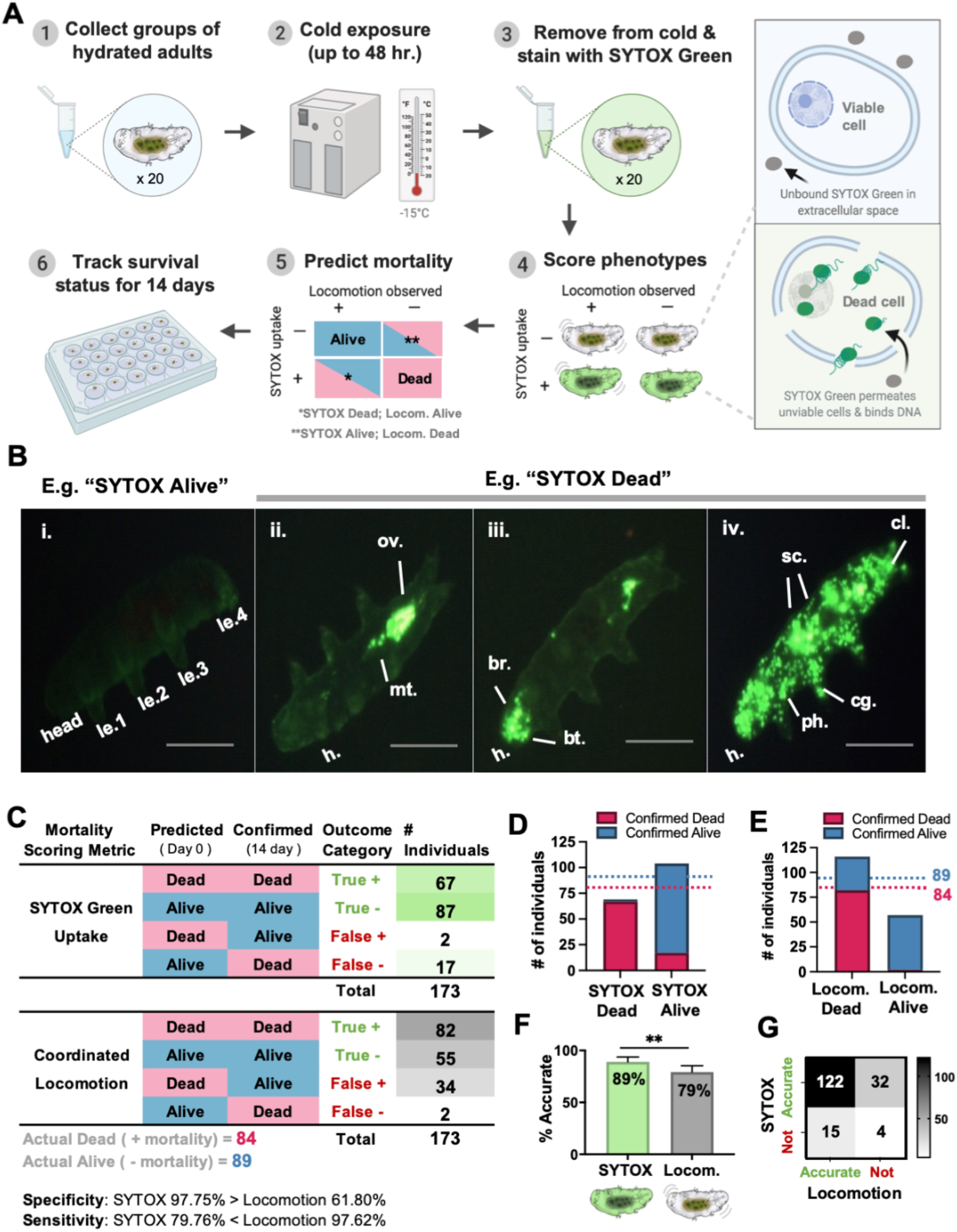
Uptake of SYTOX Green stain predicts mortality outcomes of tardigrades after cold exposure more accurately than locomotion. (A) Experimental design of the preliminary study to compare the accuracy of SYTOX Green uptake vs. the absence of locomotion in predicting mortality outcomes of cold exposed tardigrades. (B) Representative staining patterns of SYTOX Green uptake. All tardigrades are shown with heads in the lower-left corner and hind legs in the upper right, although the exact orientation varies. Le: leg, ov: ovaries, mt: malpighian tubules, br: brain, sc: storage cells, cl: cloaca. The scale bar is approximately 10 microns. Tardigrades without any staining (i) were recorded as SYTOX Alive, while those with posterior (ii), anterior and posterior (ii), or whole-body (iv) staining were recorded as SYTOX Dead. (C) Summary of predictions of long-term survival status based on SYTOX uptake and locomotion on Day 0 (“Predictions”) vs. confirmed mortality status after 14 days of observation (“Confirmed”). (D) Bar graph summarizing proportions of tardigrades predicted as dead or alive using SYTOX Green uptake, compared to confirmed values after 14 days (dotted lines: blue = 89 confirmed dead, and pink = 84 confirmed alive). (E) Bar graph summarizing proportions of tardigrades predicted as dead or alive using locomotion, compared to confirmed values after 14 days (dotted lines: blue = 89 confirmed dead, and pink = 84 confirmed alive). (F) Bar graph comparing percent accuracy (TP+TN/total) for each mortality scoring metric. SYTOX Green performed significantly better at predicting mortality according to an Exact McNemar test (p-value = 0.01315). (G) Accuracy matrix for SYTOX Green uptake and locomotion for predicting mortality, used to compute significance levels using the Exact McNemar test.

**Figure 2.**
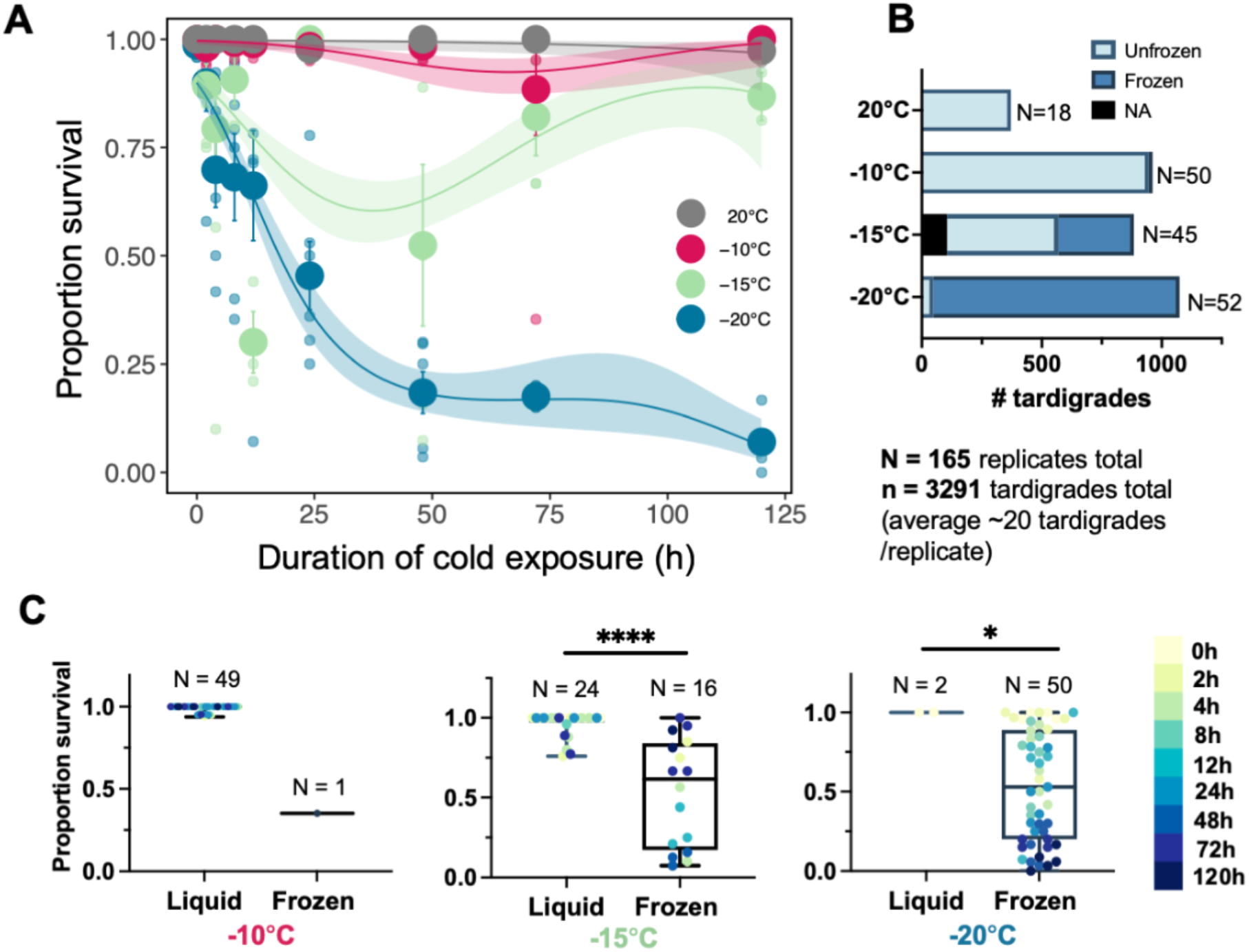
Survival of tardigrades decreases with increasing duration and intensity of cold exposure. (A) Proportion survival of tardigrades at a given intensity and duration of cold exposure. Small circles indicate the proportion of survival for each replicate (out of ∼20 individuals in a sample group), and large circles indicate the average proportion of survival between six replicates (mean ± SEM). Trendlines represent the best-fit binomial linear mixed model for that temperature (third-degree polynomial) and shading indicates 95% CI on the model predictions. Colors indicate experimental or control temperature. (B) Numbers of tardigrades among all replicates in each temperature exposure that remained liquid (light blue), froze (dark blue), or whose water status was not recorded (NA=black). N = number of sample replicates (groups of n=∼20 tardigrades in a single Eppendorf tube). The majority of samples at -10°C remained unfrozen, whereas nearly all samples froze at -20°C. The freezing status of samples at -15°C was mixed. (C) Proportion survival of samples exposed to -10°C, -15°C, or -20°C that remained liquid vs. froze. Boxes indicate median and upper and lower quartiles, with whiskers indicating the range of data. Individual data points are colored according to cold exposure time (legend right). Survival of liquid vs. frozen samples was significantly higher, at cold exposures of -15°C and -20°C (p-values=<0.0001 and p-value=0.0282, respectively, according to paired Kolmogorov-Smirnov tests).

### Cold tolerance assays interrogating duration and temperature

To establish the survival of *H. exemplaris* in response to variation in the intensity and duration of cold exposure, we exposed groups of hydrated, active adult tardigrades (n = ∼20 individuals/replicate, N = 6 replicates per treatment) to one of three ecologically-inspired cold temperatures (-10°C, -15°C -20°C) for 9 exposure times (0, 2, 4, 8, 12, 24, 48, 72 or 120h). Immediately prior to cold exposure, tardigrades were removed from standard culture conditions described above via manual microaspiration and placed into 1.5mL Eppendorf tubes in groups of 20 with 100μl of mineral water, then placed in a circulating cooler (setup described above). Once all samples were loaded into the circulating cooler, the temperature was decreased until it reached the target temperature, at which point the time 0 sample was immediately removed from the circulating cooler, and further time points were removed at 2, 4, 8, 12, 24, 48, 72 or 120h after achieving the target temperature, with 3 replicate pools per time point (performed in 2 separate batches, for a total of 6 samples, per condition). A handling control of N = 3 pools of tardigrades was treated similarly except instead of being placed in a circulating cooler was placed at room temperature (+/-2°C) in constant darkness, for the corresponding amount of time.

Immediately upon being removed from the circulating cooler, the water state of the sample (frozen or liquid) was recorded via manual visual inspection, and the tardigrades were stained in their original sample tubes with 1μM of SYTOX Green, following the protocol described in the prior section. Individuals were scored for survival based on the presence or absence of SYTOX Green uptake (using methods described above), and the proportion of survival was calculated for each pool of tardigrades (Figure 2A).

We tested the effects of cold exposure temperature and duration on survival using a generalized linear mixed model fit by maximum likelihood (Laplace Approximation) using the ‘glmerMod’ package in R (Version 4.1.3), whereby temperature, duration of the cold exposure, and their interaction were fixed effects, and replicate and batch number were random effects. Individual survival was the response variable, with a binary response of live/dead. Exposure time (duration of cold exposure) was a fixed factor, and we fit 1st, 2nd, and 3rd degree polynomials to describe the effect of exposure time, determining the best fit using AIC (delta AIC < 2 indicated the simpler model was preferred, Supplementary Figure 3). Statistical significance for the impact of factors was determined using 1st degree polynomials of tardigrade survival.

The linear mixed-effects model described continuous survival rates between 0 and 120 hours of cold exposure was plotted in ggplot2 for our various low-temperature exposures, with SEMs calculated from the 6 replicates per condition. We used this model to predict a cold hardiness metric we define as the “lethal time at low temperature” (LLt50), which is the time at which 50% of the sample population is predicted to experience mortality, at a given low temperature. We used Pearson’s Chi-squared tests with Yate’s continuity correction to determine if overall survival differed in each low temperature, compared to our 20°C control temperature.

To determine the impact of water state (frozen vs. liquid) on the survival of animals at each temperature (on average, across all time points), we conducted non-parametric Kolmogorov-Smirnov tests, calculating approximate P-values for significance between the proportion survival of liquid vs. frozen sample replicates at each temperature (GraphPad Version 9.3.1).

### Ice-inoculation assays

We next assessed how raising the freezing point of water would impact tardigrade survival of freezing, using the ice-nucleating bacteria *Pseudomonas syringae* (bacterial colony provided by Steven E. Lindow). Groups of tardigrades (n=50) were aspirated into 1.5mL Eppendorf tubes in 200μL mineral water, and inoculated with a small swab of *P. syringae* (picked from a single colony from a cultured dish, per experimental replicate) before being cooled to -10°C, -15°C, - 20°C, or a control temperature of 20°C (N=6 pools of ∼20 individuals per temperature/time combination, conducted with N=3 in 2 batches). Control tardigrades were treated identically but without the addition of *P. syringae* (N=6, of ∼20 individuals per temperature/time combination). Immediately upon thawing, the water state of each sample was recorded (frozen/unfrozen) and tardigrades were stained with SYTOX Green to score mortality as described above. Because *P. syringae* nucleates ice formation at -1.8 to -3.8°C (Maki et al. 1974), we predicted that tardigrade samples containing *P. syringae* would freeze at all subzero temperatures, whereas control samples without *P. syringae* would supercool (thus remaining liquid) to -10°C and sometimes supercooled to -15°C, based on the prior experiment. Thus, we partitioned data from this experiment into three groups “Control (no addition of *P. syringae* bacteria): Frozen,” “Control (no addition of *P. syringae* bacteria): Liquid,” and “Inoculated (with the addition of *P. syringae* bacteria): Frozen” since all cold exposed samples with *P. syringae* experienced freezing and some in the control group merely supercooled. We conducted multiple Kolmogorov-Smirnov tests, calculating approximate or exact P-values for significance between the proportion survival of sample replicates in each of the three groups at each temperature (GraphPad Version 9.3.1).

### Thermal acclimation assays

To explore if tardigrade cold tolerance may be impacted by thermal acclimation prior to cold exposure, we cultured tardigrades (as described above) in PHCBI Panasonic MIR-154-PA incubators set at 1°C, 4°C, 15°C, or on the benchtop (at 20°C) for 3 weeks. Tardigrades cultured in incubators were in constant darkness, whereas tardigrades cultured on the benchtop were in constant dim ambient light. After this 3-week acclimation period, groups of n**≈**20 tardigrades were extracted from individual culture dishes, thrice washed, and placed into 1.5mL Eppendorf tubes with 100μl of mineral water. N=3 samples were prepared from each acclimation temperature, and were then exposed to -20°C for 24 hours and 25 minutes (the time where 50% of animals died at -20°C in Figure 2). An additional N=6 samples were prepared from 20°C acclimated tardigrades (standard culture conditions, used in all prior experiments) and also exposed to -20°C for 24 hours and 25 minutes. All samples were stained with SYTOX Green post-cold exposure, as described above, and scored for tardigrade mortality rates based on the presence or absence of SYTOX Green uptake. We conducted multiple two-sided Fisher’s exact tests, calculating P-values for significance between the percent survival of each of the three temperatures compared to room temperature controls (GraphPad Version 9.3.1).

## Results

### Uptake of SYTOX Green is a more accurate measure of cold-induced mortality than lack of locomotion

To determine the best metric for scoring mortality in tardigrades post-cold exposure, we compared the accuracy of locomotion vs. SYTOX Green uptake (a live/dead cell marker) to predict long-term survival of *H. exemplaris* (n=173 animals) after a 4 to 48 hour long -15°C cold exposure (Figure 1A). Among 69/173 tardigrades that experienced uptake of SYTOX Green, we found three common staining patterns (Figure 1B): ii. Staining of the posterior (around the ovaries and malpighian tubules) iii. Staining of the anterior and posterior (head and trunk rejoin), or iv. whole-body stain (where many storage cells throughout the animal also took up stain). All three of these staining patterns were marked as SYTOX Dead (69/173 individuals), and individuals without any visible staining were marked as SYTOX Alive. We determined that 98.8% of tardigrades with any staining pattern of SYTOX Green Uptake were confirmed to have died, with SYTOX Green Uptake predicting mortality with 97.75% specificity (compared to 61.80% specificity in the case of using lack of locomotion as a marker of mortality) (Figure 1C). In the case where animals did not uptake SYTOX Green, we found that the absence of SYTOX Green uptake had a false negative rate of 2.25% but a low sensitivity of 79.76% (compared to locomotion’s sensitivity of 97.62%). Thus, SYTOX Green uptake was a highly sensitive marker of mortality, but it was a scoring metric would sometimes miss observation of lethal damage in non-stained individuals (Figure 1D). Using locomotion, however, as a marker for mortality has dramatically lower sensitivity, and would falsely classify many non-moving individuals as dead (Figure 1E). There was not a significant difference in mortality scoring accuracy in water-state specific (frozen or liquid) samples (Supplementary Figure 1), although analysis may have been limited by small sample size for partitioned data. When survival data are partitioned by exposure time, SYTOX Green performed significantly better at the longest exposure time (48 hours, Supplementary Figure 2).

Overall, we determined that SYTOX Green uptake was more accurate at predicting mortality outcomes (Accuracy = 89.01%), compared to the absence of locomotion on Day 0 (Accuracy = 79.19%.). An Exact McNemar test (with central confidence intervals) determined that SYTOX Green uptake was a significantly more accurate marker of mortality than lack of locomotion (p-value = 0.01315). These data provide evidence that SYTOX Green uptake is a more accurate metric for scoring mortality outcomes of *H. exemplaris*, and is preferable scoring methods relying on observation of locomotion after cold exposure, where organismal activity is likely to be temporarily diminished, due to sublethal damage or other physiological changes. We thus used SYTOX Green uptake as our measure of survival for subsequent experiments.

### Tardigrade mortality increases with lower subzero temperatures and longer exposure time

Temperature and exposure time significantly interacted to determine tardigrade survival (Figure 2A), when 1st degree polynomials for exposure duration were fit to survival data in a binomial generalized linear mixed model fit by maximum likelihood (Laplace Approximation), with survival based on SYTOX Green uptake as the dependent variable, duration and temperature as fixed effects, and replicate batch as a random effect (error df=2910). Both the effect of temperature, duration, and the interaction of the two factors significantly impacted survival at - 15°C vs -20°C (temperature: z-value=4.055; p-value=5.02e-05; duration: z-value=-15.507; p-value<2e-16); and the interaction of the two factors (z-value=7.925, p-value=2.27e-15) as well as between -10°C vs -20°C (temperature: z-value=9.003; p-value=<2e-16); and the interaction of the two factors (z-value=2.677; p-value=0.00742). More so, we found that overall survival at -15°C and -20°C was significantly different compared to that of our room temperature control (-20°C vs 20°C: p-value<2e-16, df=1; -15°C vs 20°C p-value<2.2e-16; df=1; Pearson’s Chi-squared test with Yates continuity correction) whereas -10°C vs 20°C was not (p-value=0.1439, df=1). Survival of room temperature controls also varied significantly with time (z-value=-13.671, p-value<2.2e-16), according to the generalized linear mixed model fit by maximum likelihood (Laplace Approximation), with duration as a fixed effect and replicate number as a random effect), although the average survival rate of the control was high at 99.50% (SEM: 0.54%).

Likewise, the survival of *H. exemplaris* was uniformly high across all durations of exposure to -10°C, with an average percent of 97.82% survival (SEM: 1.75%). The water in nearly all -10°C samples remained unfrozen at the end of each trial (Figure 2B), which we predict contributed to their high survival under these exposure conditions.

When groups of tardigrades were exposed to an intermediate temperature of -15°C, survival rates did not diminish over time in a unidirectional fashion (Figure 2A). Instead, overall survival was lowest at 12 hours (30.01% survival, SEM: 7.08%), with an average percent of survival increasing to 86.78% (SEM: 5.52%) after 120 hours of -15°C cold exposure. In the case of -15°C cold exposures, there was variation in whether a given sample supercooled or spontaneously froze, especially at shorter exposure times (less than 72 hours). We subsequently predicted that variation in tardigrade survival at -15°C may be related to the sample’s water state (frozen or liquid). Indeed, survival was significantly higher in unfrozen (liquid) samples compared to frozen samples, for tardigrades subjected to -15°C (approximate p-value <0.0001 Kolmogorov-Smirnov test) as well as for -20°C (exact p-value 0.0282 Kolmogorov-Smirnov test) (Figure 2C).

Survival of *H. exemplaris* exposed to -20°C diminished with increased exposure time (Figure 2A). An average of 98.6% (SEM: 0.88%) of animals survived after immediately reaching the low temperature of -20°C, but tardigrade survival decreased to 7.05% (SEM: 2.31%) for tardigrades exposed for the maximum duration of 120 hours. At ∼24 hours, 50% of tardigrades were dead after exposure to -20°C. We observed that the water in all but two samples exposed to -20°C froze. In Supplementary Table 5, we also observed shifts in staining patterns of SYTOX Green uptake whereby the rates of whole-body staining (compared to localized anterior or posterior staining) increased with exposure time.

In summary, these data demonstrate that *H. exemplaris* are sensitive to both intensity and duration of cold exposure, with survival decreasing precipitously with exposure time under -20°C conditions (Full data available in Supplementary Tables 2-4). Importantly, the role of ice-formation on tardigrade mortality depended on the final low temperature. In the next experiment, we thus induced ice formation at higher subzero temperatures using ice-nucleating bacteria, to explore the impact of ice formation dynamics vs. temperature alone.

### Ice-formation by ice-nucleating bacteria does not result in organismal mortality, indicating freeze tolerance

We seeded ice-nucleating bacteria *Pseudomonas syringae* into tardigrade samples and then exposed them to -10°C, -15°C, and -20°C for 120 hours and compared survival against samples without ice-nucleating bacteria, to determine the impact of controlled ice-inoculation. Without the addition of *P. syringae* into tardigrade samples, we predicted that samples exposed to -10°C would remain liquid (with high tardigrade survival), samples exposed to -20°C would freeze (with low tardigrade survival), and samples exposed to -15°C would exhibit high variation in water-state (with survival dependent on water-state)—as we observed in the large-scale survival study above (Figure 2A). Because *P. syringae* has been documented to induce ice-nucleation with high efficacy at temperatures as high as -1.8°C, we predicted that all tardigrade samples seeded with *P. syringae* before cold exposure would freeze, regardless of final low temperature (Cochet and Widehem 2000; Maki et al. 1974; Arny et al. 1976).

The addition of *P. syringae* compared to regular handling controls did not cause any observed mortality after 120h (0%) under standard culturing conditions (20°C). All cold-exposed samples seeded with *P. syringae* froze, regardless of final exposure temperature or time. Tardigrade survival was uniformly high (average 94.11%) in inoculated frozen samples across all low temperatures (Figure 3A-C). At -10°C, unfrozen controls and frozen inoculated samples had similar high survival (Figure 3B), with no statistically significant difference (p-value >0.9999, Kolmogorov-Smirnov test). At -15°C, unfrozen and frozen controls had similar high survival to inoculated frozen samples (Figure 3D), with no statistically significant difference (p-value >0.9999, Kolmogorov-Smirnov test). At -20°C, frozen controls had considerable mortality (60.99%), while survival of inoculated frozen samples remained close to 100% and was significantly higher (p-value 0.0152, Kolmogorov-Smirnov test). This illustrates that ice-formation alone does not result in tardigrade mortality, when it occurs in the presence of ice-nucleating bacteria.

**Figure 3.**
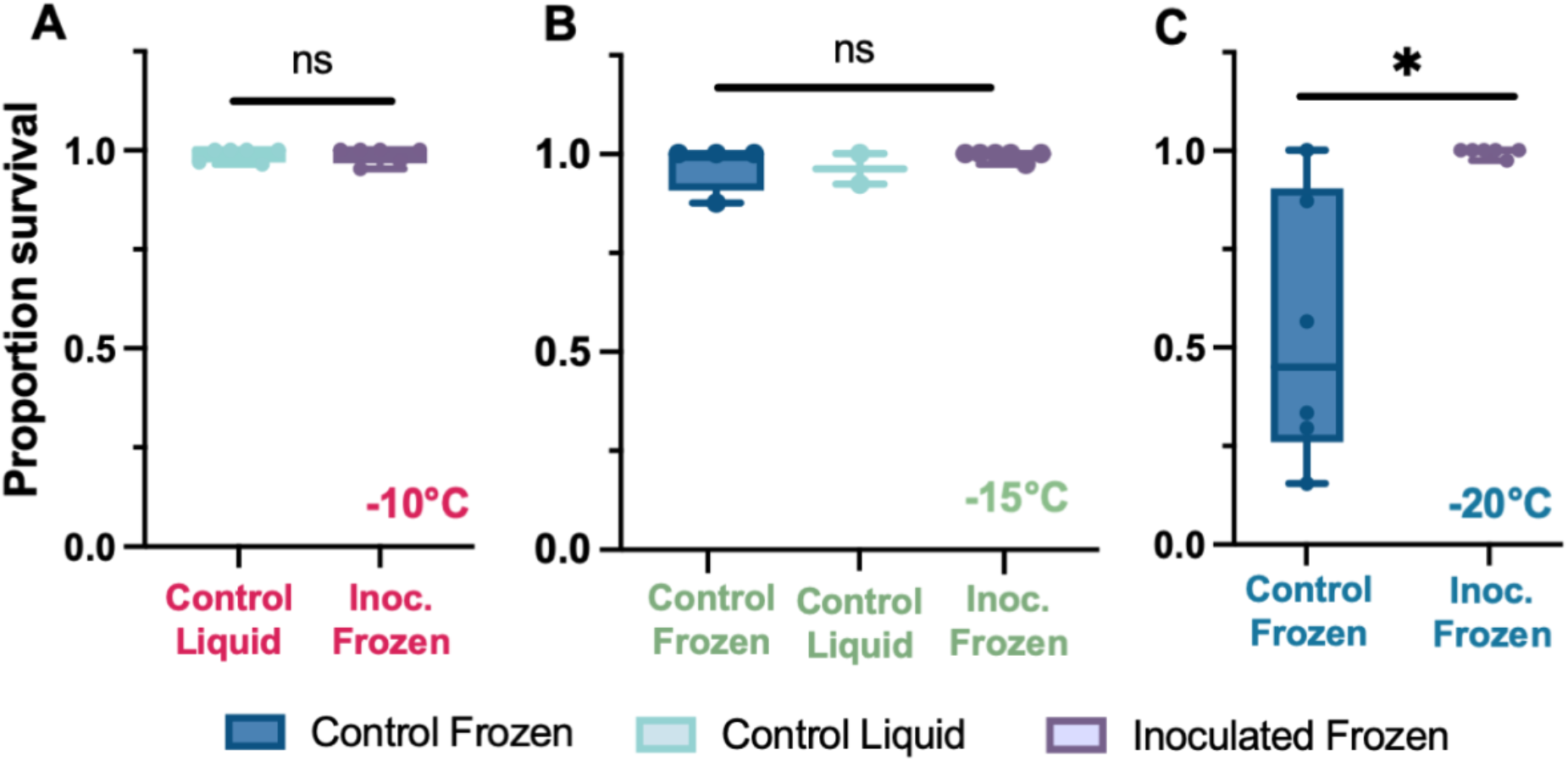
Ice-nucleating bacteria seed sample freezing but tardigrade survival remains high. (A-C) Proportion of tardigrades that survived 120 hours of cold exposure, separated by whether they received the ice-nucleating bacteria or not (Inoculated or Control) and whether the water in their tube was frozen or liquid at the end of the cold exposure (Control Frozen = dark blue, Control Liquid (unfrozen) = light blue, Inoculated Frozen = purple). Boxes indicate median and upper and lower quartiles, with whiskers indicating range of data. Individual data points represent sample replicates (tubes of ∼20 tardigrades). (A) Proportion survival of tardigrades exposed to -10°C was not significantly different among control samples that remained unfrozen and froze due to inoculation (p-value >0.9999, Kolmogorov-Smirnov test), indicating that freezing did not result in significant mortality at when induced by inoculation. (B) Proportion survival of tardigrades exposed to -15°C was not significantly different among control samples that remained unfrozen, control samples that froze, and samples that froze due to inoculation (p-value >0.9999, Kolmogorov-Smirnov tests). (C) Proportion survival of tardigrades exposed to -20°C was significantly different among control samples that froze and samples that froze due to inoculation (p-value 0.0152, Kolmogorov-Smirnov test). This indicates that the act of freezing alone does not result in mortality, but rather whether freezing is inoculated and at what temperature.

**Figure 4.**
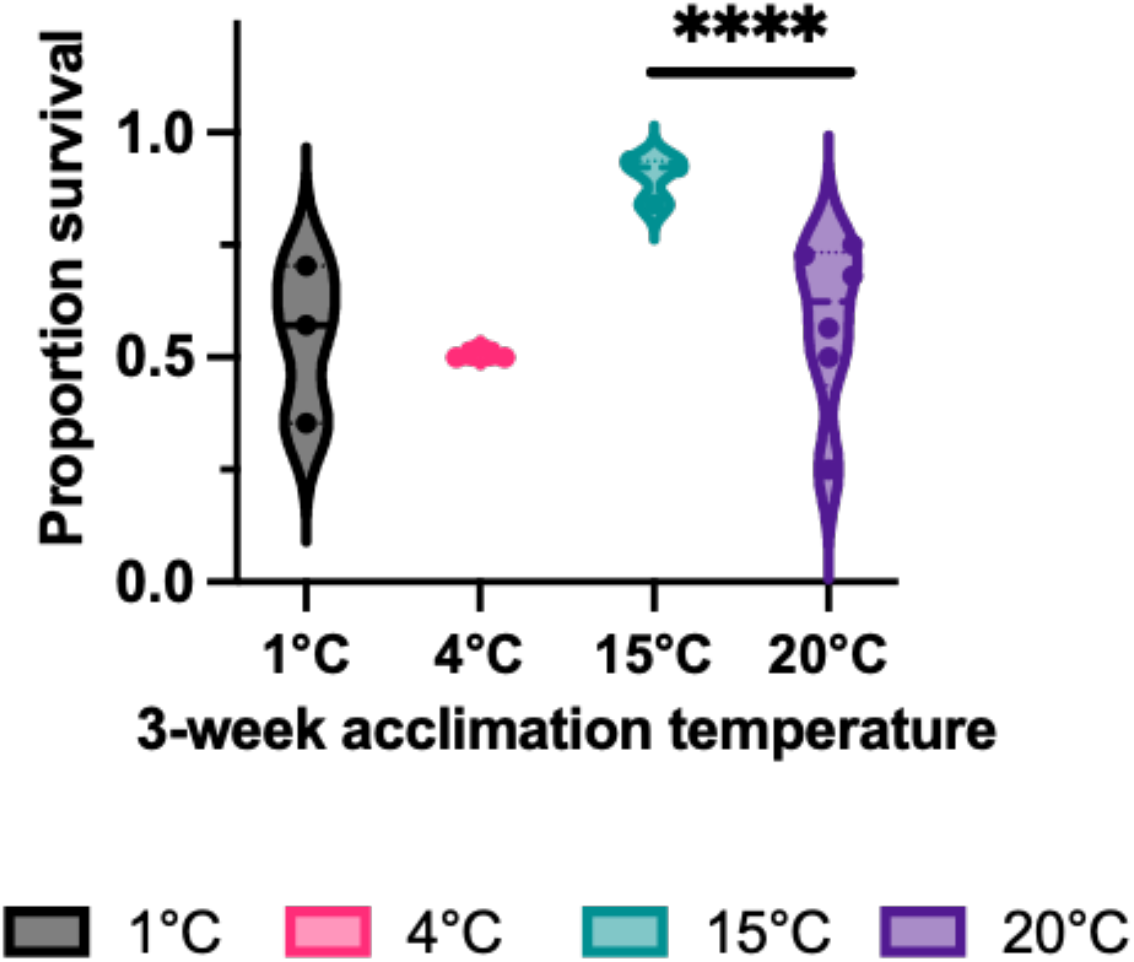
Four-week acclimation at 15°C improves tardigrade cold tolerance. Groups of tardigrades were cultured at 1°C, 4°C, 15°C or the standard 20°C culture temperature for 3 weeks and subjected to a cold exposure of -20°C for 24 hours and 25 minutes (∼50% survival in Fig 1) and scored for survival using SYTOX Green uptake. Tardigrades acclimated to 1°C and 4°C did not have significantly higher survival than 20°C control conditions (Fisher’s exact test, p=0.8868 and p=0.6709, respectively). Tardigrades acclimated to 15°C had higher survival than all other experimental thermal acclimation conditions (p= <0.0001, Fisher’s exact test). Boxes indicate median and upper and lower quartiles, with whiskers indicating the range of data.

### Three-week thermal acclimation at 15°C improves cold tolerance

Tardigrades were acclimated at three temperatures (1°C, 4°C, 15°C, 20°C) in constant darkness for 3 weeks before being exposed to -20°C for ∼24 hours (near the LLt50 for room temperature acclimated tardigrades: 24 hours and 25 minutes), to gauge the impact of thermal acclimation on cold tolerance. While 3-week acclimation at 1°C and 4°C had no significant impact on tardigrades’ survival (p-values = 0.8868 and 0.6709 respectively, Fisher’s exact test), tardigrades acclimated to 15°C had a significantly higher survival compared to room temperature controls (p-value = <0.0001, Fisher’s exact test).

## Discussion

Prior to this work, tardigrades were widely viewed as tolerant to virtually any environmental extreme, including low temperatures and freezing (Hengherr and Schill 2018). Using novel methods to score mortality, we demonstrate that survival of hydrated *Hypsibius exemplaris* is highly variable after exposure to low temperatures, depending on the interaction of duration and intensity of cold exposure, exposure to external microbial ice-nucleators, and thermal acclimation. We found that there was a threshold temperature (-20°C), at which survival declines exponentially with increasing exposure time. Above that temperature, increased exposure time did not have clear directional impacts on tardigrade survival. At this damaging low temperature, the addition of ice-nucleating bacteria to induce freezing at higher subzero temperatures dramatically increased survival, suggesting that the dynamics of ice formation and ecological conditions are critical to tardigrade cold tolerance. We established several instances where active adult tardigrades can survive external freezing of external water, without relying on anhydrobiosis. Taken in whole, we found evidence that tardigrades are not universally tolerant to environmental extremes such as cold, but given the right conditions, tardigrades can survive freezing for our longest timelines tested, without relying on tun or cyst formation. Furthermore, we provide a new framework to assay thousands of tardigrades in large-scale physiological or phenotypic screens, with improved accuracy by relying on visualization of SYTOX Green uptake.

### Cold exposure duration and intensity interact to determine tardigrade mortality

Exploring cold tolerance in hydrated, not desiccated, tardigrades is crucial to understanding the impacts of cold under more ecologically-relevant conditions. Here, we examined the survival rates of thousands of hydrated, fed, and active *H. exemplaris*. This work is the first to compare the impact of ecologically-feasible low temperature, time spent at low temperature (duration), and the interaction of these two factors on the survival of any tardigrade species. As a novel insight, we found that survival of *H. exemplaris* decreased exponentially with time when exposed to our lowest experimental temperature of -20°C (in the absence of external bacterial ice nucleators). The survival of tardigrades exposed to -20°C was high for short durations of exposure time (90.02% at the 2-hour time point), even when surrounding water froze in the tardigrade samples. Survival of tardigrades exposed to -20°C decreased, however, to an average of 45.36% after 24 hours and 7.05% after 120 hours (where all samples froze). In contrast, we found that tardigrades could tolerate temperatures of -10°C and -15°C with high survival (up to 100% depending on duration), as long as the surrounding water remained in a liquid, presumably super-cooled state. From these data alone, it appeared that *H. exemplaris* may survive low temperatures by adopting a freeze-avoidant strategy, surviving cold until exposed to a thermal threshold between -15°C and -20°C, and succumbing to freeze damage when exposed to -20°C for periods of time longer than ∼24 hours. In this work, we did not observe whether ice formed in the tissues directly, due to the methodological constraint of not being able to easily visualize tardigrades’ bodies while animals were encapsulated in opaque frozen water.

These findings are most comparable to the previously described survival rate of 55.6% for the sister species *Hypsibius dujardini*, when adult tardigrades were starved but hydrated and were cooled to -20°C for six days at a rate of 0.35°C min−1 (Guidetti et al. 2011). Guidetti et al also exposed cohorts of hydrated *H. dujardini* to final low temperatures of -9°C and -80°C, where they observed survival rates of 72.5% and 18.8% respectively (although cooling rates differed with each temperature) Overall, the authors concluded that *H. dujardini* had “moderate” cryobiotic abilities, which is consistent with our findings of *H. exemplaris* on average (with survival ranging from 7.05% to 100%, at various low temperatures and durations, in the absence of additional ice-nucleating bacteria). Our study uniquely illustrates that although *H. exemplaris* is only moderately cold-tolerant on average, it ranges from being robustly tolerant to sensitive to cold, depending on temperature and duration of cold exposure.

This work was the first study to assess the combined impacts of low temperature and duration of time spent at low temperature on tardigrade survival while holding the cooling rate roughly constant. Cooling rate has strong effects on the cold tolerance of a range of tardigrade species (Hengherr et al. 2009) and the use of a cooling circulator allowed us to control the rate of cooling during cold exposures, regardless of the final low temperature. In Guidetti *et al*, hydrated tardigrades were chilled to their target temperatures at rates that differed by 2-3x fold, which additionally impacted total time spent at low temperature and thus prevented authors from more exhaustively separating the impact of low temperature vs. duration of cold exposure (Guidetti et al. 2011). A productive avenue for future research would be to vary the rate of cooling and thawing independently of the duration and intensity of cold exposure, to allow us to assess the relative importance of each factor in determining survival.

### External ice-inoculation at higher temperatures dramatically improves survival

Initially, we found that tardigrades had higher survival of a -15°C cold exposure when animals were suspended in unfrozen water (96.1%) compared to when samples spontaneously froze (53.4%) (Figure 2C). We thus used ice-nucleating bacteria, *P. syringae*, to induce ice formation at high subzero temperatures, and to separate the effects of freezing from the effects of duration and intensity of cold exposure (Figure 3). The addition of this ice-nucleating bacteria to hydrated tardigrade samples improved tardigrade survival at the lowest exposure temperature, -20°C, significantly when frozen (99.9% compared to 58.5% for control samples; p-value >0.9999, Kolmogorov-Smirnov tests, Figure 3C). Ice-nucleation by *P. syringae* also improved survival at higher temperatures (-10°C and -15°C), where tardigrade survival averaged 97.4% and 85.1% respectively (Figure 3A & 3B). The addition of this ice-nucleating bacteria allows for ice-formation to occur at higher sub-zero temperatures, allowing a slower and more controlled rate of ice formation (Maki et al. 1974), compared to that found in tardigrade-containing samples alone. Freezing at moderate speeds (such that may occur without external ice-nucleating agents) is more likely to result in mortality, due to the predicted formation of larger ice crystals, resulting in freeze damage (Hengherr and Schill 2018).

These observations suggest that *H. exemplaris* may be tolerant of freezing when it occurs under controlled conditions of ice formation (such as at higher temperatures ranging from -1.8 to -3.8°C). This supports conclusions from several past studies, which inferred that tardigrades may experience a diversity of cold tolerance strategies including freeze avoidance, rapid cold hardening, and/or freeze tolerance, varying in accordance with the species’ natural habitat and rate of freezing (Hengherr and Schill 2018). Likewise, recent work from (Møbjerg et al. 2022) suggested that the extraordinary freeze-tolerance of *Ramazzottius varieornatus* likely relies on controlled ice formation, as suggested by observation of a significant decrease in viability following rapid cooling. However, we caution that we were not able to directly observe the freezing of the body tissues, and it is possible (though unlikely) that tardigrades were supercooled within the frozen water. Conclusively determining the cold tolerance strategy of tardigrades has remained elusive due to the destructive nature of calorimetry experiments or the methodology of scoring survival. This constraint limited the ability to discern if low temperature, external ice-formation, or ice-formation within the animal was responsible for tardigrade mortality. In future work, we will utilize micro-CT scans of cold-exposed tardigrades to visualize density shifts of intra- and extracellular freezing events.

This is the first published work to indicate that tardigrade survival may be enhanced by the presence of other cold-hardy microbes, potentially found in their shared habitats. Thus, in nature *H. exemplaris* is likely tolerant of freezing in the presence of ice inoculators. The importance of external ice nucleators is emerging as an important determinant of cold tolerance strategy in invertebrates: Even classically “freeze avoidant” species such as the Linden bug, *Pyrrhocoris apterus*, are freeze-tolerant when ice formation is inoculated externally at high sub-zero temperatures, suggesting that supercooling and freezing are not distinct strategies but rather ecophysiological alternatives, depending on the presence of external ice nucleators (Rozsypal and Košťál 2018). The role of environmental microbes on tardigrade cold tolerance and ice-formation has been absent from the literature, until data presented here.

Expanding the design of the initial study to also include examining the impact of time and temperature on the survival of tardigrades at various life history stages would be an important next step. Hengherr *et al* found no ice-nucleation events in any of the five distinct developmental stages of embryos and uniform temperatures of ice crystallization, suggesting that tardigrade embryos have similar freezing and survival patterns as adults (Hengherr et al. 2010). Yet, substantial literature in the nematode field has indicated that early-stage *C. elegans* are capable of surviving freezing via entering a daur stage, potentially by relying on the use of ice-binding proteins, demonstrating variation in cold tolerance across the life-history stages for some aquatic microorganisms (Gade et al. 2020; Kuramochi et al. 2019). Exploring whether any combination of cold exposure conditions, along with the addition of external ice-nucleating agents like *P. syringae*, shifts the survival dynamics of cold-exposed tardigrades across various life stages would better contextualize the universality of this organism’s cold tolerance.

### 15°C acclimation improves tardigrade survival rates, indicating the importance of environmental conditions for organismal growth prior to stress exposures

Few prior studies explored the impact of thermal acclimation on tardigrade stress tolerance, with the limited existing work focusing on the impact of acclimation at warmer temperatures in surviving high heat or impacting locomotor performance (Neves et al. 2020; Li and Wang 2005). Here, we found that only one of our three thermal acclimation temperatures tested (15°C) was beneficial for improving survival to sub-zero temperatures in *H. exemplaris*. This temperature of 15°C may indeed induce beneficial acclimation mechanisms prior to winter-like cold exposures, or 15°C may be a more ideal temperature to culture *H. exemplaris* for overall organismal fitness, compared to 20°C. The lack of a strong acclimation response at 1 and 4°C suggests that seasonal acclimatization might not be an important mechanism through which tardigrades survive winter cold, but further work is required to test more ecologically relevant acclimation treatments.

### SYTOX Green improves throughput and accuracy of physiological assays in tardigrades, enabling large-scale phenotypic screens

Methodologically, one of the most exciting aspects of this work was the ability to innovate on a more tractable method of scoring tardigrade mortality—via visualization of uptake of the cellular live/dead dye SYTOX Green. Virtually all prior work scored tardigrade survival by the presence of any bodily movement or locomotion, after variable amounts of time post-experiment. Our data suggest that replacing locomotion-based scoring methods with the observation of SYTOX Green uptake predicts survival-related outcomes with greater accuracy, especially in data sets such as our own, where numerous animals are initially non-mobile but subsequently recovered days or weeks after an environmental stressor. The low-false positive rate of SYTOX Green uptake as a marker of mortality will increase the utility of future physiological assays or screens, where sublethal damage (resulting in lack of initial organismal activity), or delayed onset of locomotion due to dormancy, occurs. Visualization of SYTOX Green uptake allows for mortality scoring within several hours after staining, compared to locomotion, which must be observed at precise times immediately post-cold exposure to ensure optimal accuracy and reproducibility.

In future work, SYTOX Green uptake as a metric for improved mortality scoring should be validated across a broader set of environmental stressors and, in the case of cold stress, a wider range of low temperature temperatures and exposure times. More so, additional validation studies can be conducted to ensure that SYTOX Green staining does not increase mortality in damaged animals. In Chapter 2 of this thesis, we stain tardigrades exposed to low temperatures that result in sublethal damage (-4°C for 120h, Figure 2.2) and see no significant increase in mortality compared to non-cold exposed tardigrades. While these additional findings indicate that SYTOX Green staining does not likely induce mortality in mildly damaged animals, validation studies of SYTOX Green’s accuracy and innocuous impacts on tardigrade health across a wider range of cold exposures (and thus extents of underlying damage) could better address these questions.

Nonetheless, findings here provide a pipeline to explore patterns of cold tolerance in a wide diversity of tardigrade species, in a more quantitative and high-throughput manner. This improved methodology for scoring mortality provided here serves as a foundation to further automate phenotypic screens in tardigrades, which could rely on automated imaging collection techniques seen in other microscopic invertebrates or cells, rather than relying on manual inspection alone (Pulak 2006; Dravid et al. 2021). The majority of prior studies examined the survival of dozens to hundreds of tardigrades (Hengherr and Schill 2018), but these improved methods enable the study of thousands of individuals within a single experiment, enabling a higher standard for robust sample sizes in future work.

## Conclusions

By relying on visualization of SYTOX Green uptake to score tardigrade mortality, our work examined the effects of several ecologically-relevant environmental factors on cold tolerance at large-scale, with improved accuracy and speed, compared to prior studies that rely on scoring mortality based on locomotion alone. In contrast to the popular science narrative of tardigrades as universally tolerant to all environmental extremes, we found that the survival of *Hypsibius exemplaris* is significantly decreased with lower subzero temperatures and prolonged exposure times, unless animals are frozen alongside external ice-nucleating agents like the bacteria *P. syringae* or prior thermal acclimation at 15°C. This work challenges the narrative that tardigrades are universally tolerant to all environmental stressors, and furthermore demonstrates a more nuanced view of how more ecologically relevant conditions impact tardigrade survival and performance. Our work emphasizes the importance of external ice nucleators on cold tolerance and suggests that bacteria present in the environment of tardigrades likely improve cold tolerance markedly over survival measured in pure water in the laboratory. This illustrates the importance of considering external ice nucleators more generally in invertebrate cold tolerance strategies.

This work provides key scaffolding for other large-scale phenotypic assays that rely on scoring survival in tardigrades, across a wide range of environmental stressors. More so, we provide a framework for studying stress tolerance across a wide range of tardigrade species, enabling researchers to explore tardigrade physiology through a comparative lens with improved throughput. These advancements further solidify tardigrades as an increasingly tractable emerging model organism, across a growing set of biological disciplines.

## Supporting information

All supplemental figures

## Acknowledgements

Our work was partially supported by several Walker Grants from the Essig Museum of Entomology (2017-2021), Summer Research Awards from the Department of Integrative Biology, UC Berkeley (2017-2022), and an IB Dissertation Completion Award (2021). Kevin T. Roberts performed GLM modeling and consulted in data analysis. Daisy Stock and Kylie Cheng aided in tardigrade viability in Figure 1. Déjenaé See, Kylie Cheng, Lily Shang, and Daisy Stock assisted in tardigrade husbandry. Steven Lindow generously provided *P. syringae* cultures. Michael Shapira generously provided use of fluorescent microscopy equipment. Caroline Williams provided helpful comments in crafting and editing this manuscript.

