## Supplementary material for "Survival of tardigrades (*Hypsibius exemplaris*) to subzero temperatures depends on exposure intensity, duration, and ice-nucleation—as shown by large-scale mortality dye-based assays": All supplemental figures

**A** FROZEN SAMPLES

i

| Mortality Scoring Metric | Predicted ( Day 0 ) | Confirmed (14 day ) | Outcome Category | # Individuals |
|---|---|---|---|---|
| SYTOX Green Uptake | Dead | Dead | True + | 61 |
| | Alive | Alive | True - | 26 |
| | Dead | Alive | False + | 1 |
| | Alive | Dead | False - | 8 |
| | | | Total | 96 |
| Coordinated Locomotion | Dead | Dead | True + | 68 |
| | Alive | Alive | True - | 13 |
| | Dead | Alive | False + | 14 |
| | Alive | Dead | False - | 1 |
| | | | Total | 96 |

Actual Dead ( + mortality) = 69
Actual Alive ( - mortality) = 27

**Specificity**: SYTOX 92.30% > Locomotion 48.15%
**Sensitivity**: SYTOX 88.41% < Locomotion 98.56%

ii

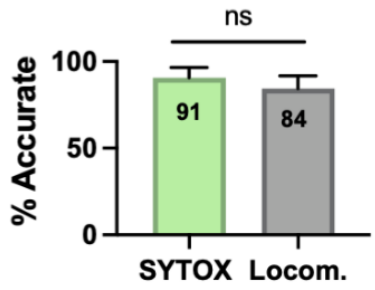


iii

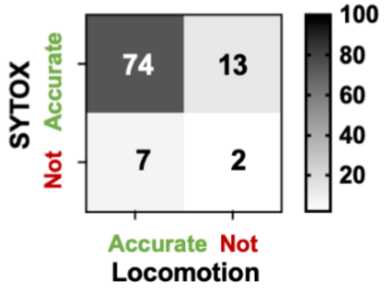


**B**

i NOT FROZEN SAMPLES

| Mortality Scoring Metric | Predicted ( Day 0 ) | Confirmed (14 day ) | Outcome Category | # Individuals |
|---|---|---|---|---|
| SYTOX Green Uptake | Dead | Dead | True + | 6 |
| | Alive | Alive | True - | 61 |
| | Dead | Alive | False + | 1 |
| | Alive | Dead | False - | 9 |
| | | | Total | 77 |
| Coordinated Locomotion | Dead | Dead | True + | 14 |
| | Alive | Alive | True - | 42 |
| | Dead | Alive | False + | 20 |
| | Alive | Dead | False - | 1 |
| | | | Total | 77 |

Actual Dead ( + mortality) = 15
Actual Alive ( - mortality) = 62

**Specificity**: SYTOX 98.39% > Locomotion 67.74%
**Sensitivity**: SYTOX 40% < Locomotion 93.33%

ii

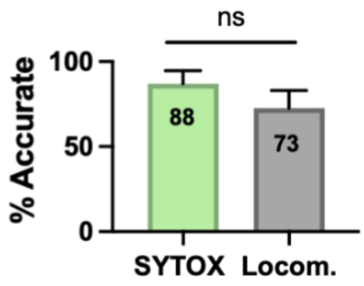


iii

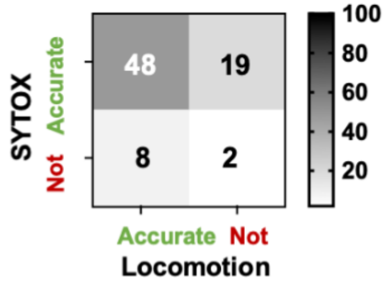


**Supplementary Figure 1. Accuracy of mortality scoring metrics by water-state (frozen vs. not frozen)**

**Supplementary Figure 1. continued**

(A) i. Table summarizing predictions of long-term survival status based on SYTOX uptake and locomotion on Day 0 (“Predictions”) vs. confirmed mortality status after 14 days of observation (“Confirmed”), for samples that froze in Fig 1.

ii. Bar graph comparing percent accuracy (TP+TN/total) for each mortality scoring metric, for frozen samples from Figure 1. SYTOX Green did not perform significantly better at predicting mortality according to an Exact McNemar test (p-value = 0.2632).

iii. Accuracy matrix for SYTOX Green uptake and locomotion for predicting mortality in frozen samples, used to compute significance levels using the Exact McNemar test.

(B) i. Table summarizing predictions of long-term survival status based on SYTOX uptake and locomotion on Day 0 (“Predictions”) vs. confirmed mortality status after 14 days of observation (“Confirmed”), for samples that remained liquid in Fig 1.

ii. Bar graph comparing percent accuracy (TP+TN/total) for each mortality scoring metric, for liquid samples from Figure 1. SYTOX Green did not perform significantly better at predicting mortality according to an Exact McNemar test (p-value = 0.05224).

iii. Accuracy matrix for SYTOX Green uptake and locomotion for predicting mortality in liquid samples, used to compute significance levels using the Exact McNemar test.

**A**

| Mortality Scoring Metric | Predicted ( Day 0 ) | Confirmed (14 day ) | Outcome Category | # Individuals 4 hours | # Individuals 8 hours | # Individuals 12 hours | # Individuals 48 hours |
|---|---|---|---|---|---|---|---|
| SYTOX Green Uptake | Dead | Dead | True + | 11 (18) | 5 (13) | 28 (30) | 23 (23) |
| | Alive | Alive | True - | 24 (25) | 30 (30) | 8 (9) | 25 (25) |
| | Dead | Alive | False + | 1 | 0 | 1 | 0 |
| | Alive | Dead | False - | 7 | 8 | 2 | 0 |
| | | | Total | 43 | 43 | 3 | 48 |
| Coordinated Locomotion | Dead | Dead | True + | 17 (18) | 12 (13) | 30 (30) | 23 (23) |
| | Alive | Alive | True - | 22 (25) | 11 (30) | 8 (9) | 14 (25) |
| | Dead | Alive | False + | 3 | 19 | 1 | 11 |
| | Alive | Dead | False - | 1 | 1 | 0 | 0 |
| Actual Dead ( + mortality) | | | Total | 43 | 43 | 39 | 48 |
| Actual Alive ( - mortality) | | | | | | | |

**B**

| SYTOX Green Uptake | 4 hours | 8 hours | 12 hours | 48 hours |
|---|---|---|---|---|
| % Accuracy | **81.4** | **81.4** | **92.31** | **100** |
| % Specificity | 96 | 100 | 88.89 | 100 |
| % Sensitivity | 61.11 | 38.46 | 93.33 | 100 |

| Coordinated Locomotion | 4 hours | 8 hours | 12 hours | 48 hours |
|---|---|---|---|---|
| % Accuracy | **90.7** | **53.49** | **97.44** | **77.08** |
| % Specificity | 88 | 36.67 | 88.89 | 56 |
| % Sensitivity | 94.44 | 92.31 | 100 | 100 |

Exact McNemar test (SYTOX vs. Locomotion accuracy matrix)

| | 4 hours | 8 hours | 12 hours | 48 hours |
|---|---|---|---|---|
| p-values | 0.2891 | 0.02896 | 0.25 | 0.0009766 |
| Significance | NS | * | NS | *** |

**C**

| 4 hours | Locomotion | |
|---|---|---|
| Sytox | Accurate | Inaccurate |
| Accurate | 33 | 6 |
| Inaccurate | 2 | 2 |

| 8 hours | Locomotion | |
|---|---|---|
| Sytox | Accurate | Inaccurate |
| Accurate | 16 | 19 |
| Inaccurate | 7 | 1 |

| 12 hours | Locomotion | |
|---|---|---|
| Sytox | Accurate | Inaccurate |
| Accurate | 36 | 0 |
| Inaccurate | 2 | 1 |

| 48 hours | Locomotion | |
|---|---|---|
| Sytox | Accurate | Inaccurate |
| Accurate | 37 | 0 |
| Inaccurate | 11 | 0 |

**Supplementary Figure 2. Accuracy of mortality scoring metrics across timepoints (regardless of freezing status)**

(A) Table summarizing predictions of long-term survival status based on SYTOX Green uptake and locomotion on Day 0 (“Predictions”) vs. confirmed mortality status after 14 days of observation (“Confirmed”), for samples in Fig 1. according to -15°C exposure time (4-48 h).

(B) Tables summarizing accuracy metrics and p-values (Exact McNemar tests) of SYTOX Green uptake vs. locomotion, for samples in Fig 1. according to -15°C exposure time (4-48 h).

(C) Accuracy matrix for SYTOX Green uptake and locomotion for predicting mortality in time course-separated samples, used to compute significance levels using the Exact McNemar test.

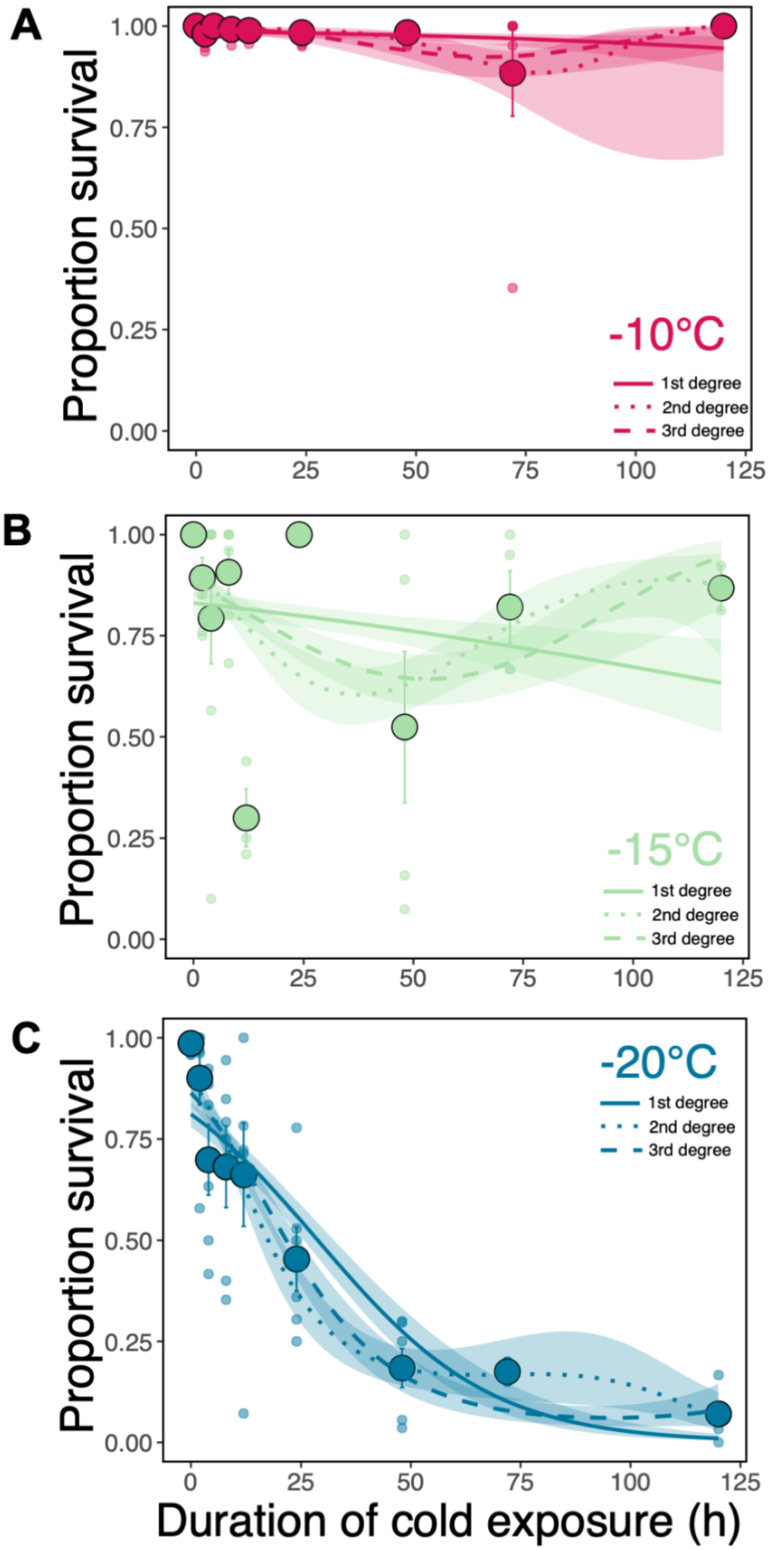


**Supplementary Figure 3. Selection of best-fit for proportion survival across timepoints for general linear model**

**Supplementary Figure 3. continued**
(A) Binomial polynomial fit for the -10°C treatments for first, second, and third-degree polynomials. Best-fit based on AIC is third-degree polynomial (AIC=171.5718) vs AIC values for second-degree (AIC=175.837) or first-degree polynomials (AIC=192.7941). Second-degree polynomial is the best fit for -10°C survival data according to BIC (190.444), compared to first- and third-degree (BIC=202.5322 and 191.0479, respectively).
(B) Binomial polynomial fit for the -15°C treatments for first, second, and third-degree polynomials. Best-fit based on AIC is third-degree polynomial (AIC=845.1951) vs AIC values for second-degree (AIC=853.8989) or first-degree polynomials (AIC=884.2961). Third-degree polynomial is the best fit for -15°C survival data according to BIC (864.333), compared to first- and second-degree (BIC=893.8651 and 868.2523, respectively).
(C) Binomial polynomial fit for the -20°C treatments for first, second, and third-degree polynomials. Best-fit based on AIC is third-degree polynomial (AIC=1017.132) vs AIC values for second-degree (AIC=1028.914) or first-degree polynomials (AIC=1067.566). Third-degree polynomial is the best fit for -20°C survival data according to BIC (1037.049), compared to first- and third-degree (BIC=1043.851 and 1077.524, respectively).

**Proportion Survival (non-repeated measurements)**

| | Replicate | 0 hrs. | 2 hrs. | 4 hrs. | 8 hrs. | 12 hrs. | 24 hrs. | 48 hrs. | 72 hrs. | 120 hrs. |
|---|---|---|---|---|---|---|---|---|---|---|
| | 1 | 1 | 1 | 0.63 | 0.35 | 0.71 | 0.3 | 0.3 | 0.15 | 0.09 |
| **–20°C** | 2 | 1 | 0.58 | 0.42 | 0.4 | 1 | 0.36 | 0.17 | 0.2 | 0.03 |
| **Survival** | 3 | 1 | 1 | 0.83 | 0.85 | 0.68 | 0.53 | 0.06 | 0.15 | 0 |
| **among reps.** | 4 | 1 | 0.89 | 0.5 | 0.94 | 0.78 | 0.25 | 0.04 | 0.2 | 0.17 |
| | 5 | 0.96 | 0.97 | 0.88 | 0.79 | 0.72 | 0.5 | 0.3 | NA | 0.06 |
| | 6 | 0.96 | 0.96 | 0.92 | 0.75 | 0.07 | 0.78 | 0.25 | NA | NA |
| **Overall average by time** | | **0.99** | **0.9** | **0.7** | **0.68** | **0.66** | **0.45** | **0.19** | **0.18** | **0.07** |
| Liquid average by time | | 1 | NA | NA | NA | NA | NA | NA | NA | NA |
| Frozen average by time | | **0.98** | **0.94** | **0.77** | **0.8** | **0.58** | **0.5** | **0.17** | **0.18** | **0.08** |

| | |
|---|---|
| Overall **liquid** average | **1** |
| Overall **frozen** average | **0.54** |

**# of Tardigrades Scored**

| | Replicate | 0 hrs. | 2 hrs. | 4 hrs. | 8 hrs. | 12 hrs. | 24 hrs. | 48 hrs. | 72 hrs. | 120 hrs. |
|---|---|---|---|---|---|---|---|---|---|---|
| | 1 | 25 | 6 | 30 | 17 | 21 | 23 | 27 | 20 | 23 |
| **–20°C** | 2 | 22 | 19 | 12 | 25 | 18 | 25 | 24 | 15 | 30 |
| **# Tardigrades** | 3 | 11 | 18 | 30 | 33 | 25 | 17 | 18 | 20 | 18 |
| **among reps.** | 4 | 18 | 28 | 32 | 18 | 23 | 16 | 28 | 15 | 12 |
| | 5 | 24 | 31 | 26 | 24 | 18 | 18 | 10 | NA | 17 |
| | 6 | 24 | 26 | 26 | 24 | 14 | 9 | 8 | NA | NA |
| **Total by time** | | **124** | **128** | **156** | **141** | **119** | **108** | **115** | **70** | **100** |
| **Overall total, all times** | | **1061** | | | | | | | | |

**Supplementary Table 1. Summary of survival data and # of tardigrades used for -20°C samples** Light blue indicates samples that froze, whereas Green or red gradients represent variation in the proportion of survival or the number of tardigrades used in each sample, respectively. Grey NA boxes indicate experimental conditions that were not measured, due to human or instrument error.

**Proportion Survival (non-repeated measurements)**

| | Replicate | 0 hrs. | 2 hrs. | 4 hrs. | 8 hrs. | 12 hrs. | 24 hrs. | 48 hrs. | 72 hrs. | 120 hrs. |
|---|---|---|---|---|---|---|---|---|---|---|
| | 1 | 1 | 1 | 1 | 0.96 | 0.44 | 1 | 0.89 | 0.67 | 0.81 |
| **–15°C** | 2 | 1 | 0.76 | 1 | 1 | 0.25 | 1 | 0.77 | 0.67 | 0.92 |
| **Survival** | 3 | 1 | 1 | 1 | 1 | 0.21 | 1 | 1 | 0.95 | NA |
| **among reps.** | 4 | 1 | 1 | 1 | 0.68182 | NA | 1 | 0.16 | 1 | NA |
| | 5 | 1 | 0.85 | 0.8 | 0.8 | NA | NA | 0.13 | NA | NA |
| | 6 | 1 | 0.75 | 0.88 | 1 | NA | NA | 0.07 | NA | NA |
| | 7 | | | 0.1 | | | | | | |
| | 8 | | | 0.57 | | | | | | |
| **Overall average by time** | | **1** | **0.89** | **0.79** | **0.91** | **0.3** | **1** | **0.5** | **0.82** | **0.87** |
| Liquid average by time | | 1 | 0.94 | 0.95 | 0.96 | NA | 1 | 0.89 | NA | NA |
| Frozen average by time | | | 0.8 | 0.34 | NA | 0.3 | NA | 0.12 | NA | NA |

| | |
|---|---|
| Overall **liquid** average | **1** |
| Overall **frozen** average | **0.79** |
| Overall unknown avg. | **0.9** |

**# of Tardigrades Scored**

| | Replicate | 0 hrs. | 2 hrs. | 4 hrs. | 8 hrs. | 12 hrs. | 24 hrs. | 48 hrs. | 72 hrs. | 120 hrs. |
|---|---|---|---|---|---|---|---|---|---|---|
| | 1 | 25 | 15 | 20 | 25 | 25 | 19 | 18 | 24 | 16 |
| **–15°C** | 2 | 16 | 25 | 20 | 25 | 24 | 25 | 22 | 18 | 13 |
| **# Tardigrades** | 3 | 20 | 16 | 18 | 10 | 19 | 19 | 27 | 20 | NA |
| **among reps.** | 4 | 15 | 16 | 19 | 22 | NA | 21 | 19 | 24 | NA |
| | 5 | 12 | 20 | 15 | 25 | NA | NA | 16 | NA | NA |
| | 6 | 12 | 16 | 25 | 23 | NA | NA | 27 | NA | NA |
| | 7 | | | 10 | | | | | | |
| | 8 | | | 23 | | | | | | |
| | **Total** | **100** | **108** | **150** | **130** | **68** | **84** | **129** | **86** | **29** |
| **Overall total, all times** | | **884** | | | | | | | | |

**Supplementary Table 2. Summary of survival data and # of tardigrades used for -15°C samples** Light blue indicates samples that froze, whereas green or red gradients represent variation in the proportion of survival or the number of tardigrades used in each sample, respectively. Grey NA boxes indicate experimental conditions that were not measured, due to human or instrument error.

**Proportion Survival (non-repeated measurements)**

| | Replicate | 0 hrs. | 2 hrs. | 4 hrs. | 8 hrs. | 12 hrs. | 24 hrs. | 48 hrs. | 72 hrs. | 120 hrs. |
|---|---|---|---|---|---|---|---|---|---|---|
| | 1 | 1 | 1 | 1 | 0.95 | 0.96 | 0.95 | 0.95 | 0.35 | 1 |
| –10°C | 2 | 1 | 1 | 1 | 1 | 1 | 1 | 0.95 | 1 | 1 |
| Survival | 3 | 1 | 0.95 | 1 | 1 | 1 | 0.95 | 1 | 1 | 1 |
| among reps. | 4 | 1 | 0.94 | 1 | 1 | 1 | 1 | 1 | 0.95 | 1 |
| | 5 | 1 | 1 | 1 | 1 | NA | 1 | 1 | 1 | NA |
| | 6 | 1 | 1 | 1 | 1 | NA | 1 | 1 | 1 | NA |
| **Overall average by time** | | **1** | **0.98** | **1** | **0.99** | **0.99** | **0.98** | **0.98** | **0.88** | **1** |
| Liquid average by time | | 1 | 0.98 | 1 | 0.99 | 0.99 | 0.98 | 0.98 | 0.99 | 1 |
| Frozen average by time | | NA | NA | NA | NA | NA | NA | NA | 0.35 | NA |

| | |
|---|---|
| Overall **liquid** average | **0.99** |
| Overall **frozen** average | **0.98** |

**# of Tardigrades Scored**

| | Replicate | 0 hrs. | 2 hrs. | 4 hrs. | 8 hrs. | 12 hrs. | 24 hrs. | 48 hrs. | 72 hrs. | 120 hrs. |
|---|---|---|---|---|---|---|---|---|---|---|
| | 1 | 23 | 22 | 21 | 21 | 23 | 20 | 20 | 17 | 18 |
| –10°C | 2 | 23 | 17 | 19 | 21 | 19 | 18 | 22 | 19 | 23 |
| # Tardigrades | 3 | 26 | 19 | 16 | 18 | 22 | 22 | 17 | 21 | 27 |
| among reps. | 4 | 21 | 16 | 17 | 21 | 19 | 16 | 15 | 21 | 25 |
| | 5 | 19 | 18 | 16 | 19 | NA | 21 | 16 | 10 | NA |
| | 6 | 15 | 16 | 19 | 25 | NA | 12 | 17 | 14 | NA |
| **Total by time** | | **127** | **108** | **108** | **125** | **83** | **109** | **107** | **102** | **93** |
| **Overall total, all times** | | **962** | | | | | | | | |

**Supplementary Table 3. Summary of survival data and # of tardigrades used for -10°C samples** Light blue indicates samples that froze, whereas green or red gradients represent variation in the proportion of survival or the number of tardigrades used in each sample, respectively. Grey NA boxes indicate experimental conditions that were not measured, due to human or instrument error.

**Proportion Survival (non-repeated measurements)**

| | Replicate | 0 hrs. | 2 hrs. | 4 hrs. | 8 hrs. | 12 hrs. | 24 hrs. | 48 hrs. | 72 hrs. | 120 hrs. |
|---|---|---|---|---|---|---|---|---|---|---|
| 20°C | 1 | 1 | 1 | 1 | 1 | 1 | 0.95 | 1 | 1 | 0.95 |
| Survival | 2 | 1 | 1 | 1 | 1 | 1 | 1 | 1 | 1 | 1 |
| **Overall average by time** | | **1** | **1** | **1** | **1** | **1** | **0.98** | **1** | **1** | **0.98** |
| Liquid average by time | | 1 | 1 | 1 | 1 | 1 | 0.98 | 1 | 1 | 0.98 |
| Frozen average by time | | NA | NA | NA | NA | NA | NA | NA | 0.35 | NA |

| | |
|---|---|
| Overall **liquid** average | **0.99** |
| Overall **frozen** average | **NA** |

**# of Tardigrades Scored**

| | Replicate | 0 hrs. | 2 hrs. | 4 hrs. | 8 hrs. | 12 hrs. | 24 hrs. | 48 hrs. | 72 hrs. | 120 hrs. |
|---|---|---|---|---|---|---|---|---|---|---|
| 20°C | 1 | 21 | 18 | 28 | 21 | 20 | 22 | 22 | 15 | 38 |
| # Tardigrades | 2 | 21 | 19 | 21 | 8 | 20 | 19 | 22 | 16 | 30 |
| **Total by time** | | **42** | **37** | **49** | **29** | **40** | **41** | **44** | **31** | **68** |
| **Overall total, all times** | | **381** | | | | | | | | |

**Supplementary Table 4. Summary of survival data and # of tardigrades used for 20°C control samples** Light blue indicates samples that froze, whereas green or red gradients represent variation in the proportion of survival or the number of tardigrades used in each sample, respectively. Grey NA boxes indicate experimental conditions that were not measured, due to human or instrument error.

**–20°C**

| SYTOX Uptake | Status | 0 hrs. | 2 hrs. | 4 hrs. | 8 hrs. | 12 hrs. | 24 hrs. | 48 hrs. | 72 hrs. | 120 hrs. | Total |
|---|---|---|---|---|---|---|---|---|---|---|---|
| | | # of Tardigrades Scored | | | | | | | | | |
| None | Alive | 122 | 115 | 112 | 98 | 90 | 44 | 19 | 12 | 9 | 621 |
| Anterior | Dead | 0 | 0 | 1 | 1 | 0 | 0 | 2 | 0 | 0 | 4 |
| Posterior | Dead | 1 | 2 | 4 | 4 | 7 | 7 | 14 | 2 | 20 | 61 |
| Ant.-Post. | Dead | 1 | 5 | 2 | 0 | 0 | 3 | 8 | 0 | 10 | 29 |
| Whole-body | Dead | 0 | 6 | 37 | 35 | 21 | 52 | 67 | 56 | 72 | 346 |
| | Total | **124** | **128** | **156** | **138** | **118** | **106** | **110** | **70** | **111** | **1061** |

**–15°C**

| SYTOX Uptake | Status | 0 hrs. | 2 hrs. | 4 hrs. | 8 hrs. | 12 hrs. | 24 hrs. | 48 hrs. | 72 hrs. | 120 hrs. | Total |
|---|---|---|---|---|---|---|---|---|---|---|---|
| | | # of Tardigrades Scored | | | | | | | | | |
| None | Alive | 100 | 94 | 125 | 117 | 20 | 84 | 69 | 71 | 25 | 705 |
| Anterior | Dead | 0 | 2 | 0 | 1 | 2 | 0 | 0 | 0 | 0 | 5 |
| Posterior | Dead | 0 | 4 | 3 | 1 | 0 | 0 | 20 | 15 | 1 | 44 |
| Ant.-Post. | Dead | 0 | 0 | 0 | 0 | 0 | 0 | 0 | 0 | 0 | 0 |
| Whole-body | Dead | 0 | 8 | 22 | 11 | 46 | 0 | 40 | 0 | 3 | 130 |
| | Total | **100** | **108** | **150** | **130** | **68** | **84** | **129** | **86** | **29** | **884** |

**–10°C**

| SYTOX Uptake | Status | 0 hrs. | 2 hrs. | 4 hrs. | 8 hrs. | 12 hrs. | 24 hrs. | 48 hrs. | 72 hrs. | 120 hrs. | Total |
|---|---|---|---|---|---|---|---|---|---|---|---|
| | | # of Tardigrades Scored | | | | | | | | | |
| None | Alive | 127 | 106 | 108 | 123 | 82 | 107 | 107 | 90 | 93 | 943 |
| Anterior | Dead | 0 | 0 | 0 | 0 | 0 | 0 | 0 | 0 | 0 | 0 |
| Posterior | Dead | 0 | 0 | 0 | 0 | 0 | 1 | 0 | 7 | 0 | 8 |
| Ant.-Post. | Dead | 0 | 0 | 0 | 0 | 0 | 0 | 0 | 0 | 0 | 0 |
| Whole-body | Dead | 0 | 2 | 0 | 2 | 1 | 1 | 0 | 5 | 0 | 11 |
| | Total | **127** | **108** | **108** | **125** | **83** | **109** | **107** | **102** | **93** | **962** |

**Supplementary Table 5. Summary of # of tardigrades exhibiting various SYTOX Green uptake patterns, across each cold exposure's timecourse** Green gradients represent variation in the number of tardigrades that did not uptake any SYTOX Green stain (and were scored to survive). Red gradients represent variation in the number of tardigrades that exhibited one of the four SYTOX Green staining patterns (and were scored to have died), according to those shown in Figure 1B.
